# FicD regulates adaptation to the unfolded protein response in the murine liver

**DOI:** 10.1101/2024.04.15.589620

**Authors:** Amanda K. Casey, Nathan M Stewart, Naqi Zaidi, Hillery F. Gray, Amelia Cox, Hazel A. Fields, Kim Orth

## Abstract

The unfolded protein response (UPR) is a cellular stress response that is activated when misfolded proteins accumulate in the endoplasmic reticulum (ER). The UPR elicits a signaling cascade that results in an upregulation of protein folding machinery and cell survival signals. However, prolonged UPR responses can result in elevated cellular inflammation, damage, and even cell death. Thus, regulation of the UPR response must be tuned to the needs of the cell, sensitive enough to respond to the stress but pliable enough to be stopped after the crisis has passed. Previously, we discovered that the bi-functional enzyme FicD can modulate the UPR response via post-translational modification of BiP. FicD AMPylates BiP during homeostasis and deAMPylates BiP during stress. We found this activity is important for the physiological regulation of the exocrine pancreas. Here, we explore the role of FicD in the murine liver. Like our previous studies, livers lacking FicD exhibit enhanced UPR signaling in response to short term physiologic fasting and feeding stress. However, the livers of *FicD*^−/−^ did not show marked changes in UPR signaling or damage after either chronic high fat diet (HFD) feeding or acute pathological UPR induction. Intriguingly, *FicD*^−/−^ mice showed changes in UPR induction and weight loss patterns following repeated pathological UPR induction. These findings show that FicD regulates UPR responses during mild physiological stress and may play a role in maintaining resiliency of tissue through adaptation to repeated ER stress.

## Introduction

Cells must maintain a careful balance of protein synthesis, modification, and degradation to sustain homeostasis despite the changing environmental and functional demands of an organism. When this balance is disrupted, cells respond with cellular stress alarms to activate complex signaling networks that can promote both cell survival and cell death pathways. Tissues with fluctuating physiological demands rely on cellular stress pathways to maintain cellular homeostasis and protect cell survival. The unfolded protein response (UPR) is a cellular stress response that plays a critical role in the regulation of protein homeostasis and is triggered through a variety of stresses in the endoplasmic reticulum (ER). Tuning the UPR for each cell type is needed for healthy regulation of normal tissue function during ER stress as disruption of UPR signaling contributes to a variety of diseases in a broad range of tissues (1–10).

Activation of the UPR is regulated by the ER chaperone protein BiP (also known as Grp78 or Hspa5), which leads to a complex signaling network that promotes both cell survival and apoptotic pathways (11, 12). The UPR consists of three distinct intertwined signaling branches, controlled by the three UPR transducers: Inositol-requiring enzyme-1α (IRE1), protein kinase R-like endoplasmic reticulum kinase (PERK), and activating transcription factor 6 (ATF6). These three branches are activated when unfolded proteins accumulate in the lumen of the ER and bind to BiP, resulting in the disassociation of BiP from the three transducers (13). Together, their activation bolsters the protein folding capacity of the ER through specific translation of protein quality control machinery with concurrent repression of global protein translation. Activation of these UPR transducers also leads to stimulation of metabolic, pro-inflammatory, and apoptotic signaling pathways (14–17).

The UPR has evolved to function beyond the role of a simple stress response pathway. Mounting evidence supports the UPR serving as a tool of adaptation that can sense and incorporate information to proactively buffer anticipated challenges to cellular homeostasis in a tissue specific manner (18–22). Rigorous prior research studies indicate that the three branches of the UPR can be activated independently depending on the cell type and the cause and level of stress, leading to changes in coordination of signaling (23). The importance of these tissue-specific differences in UPR signaling is evident as many individual tissues and organs rely on a subset of the UPR signaling molecules for proper development and function (16, 17, 24–26). However, the mechanisms that determine a specific response remain unclear.

FicD (known as Fic or HYPE), is an ER resident enzyme conserved across all metazoans(27). This enzyme is responsible for the reversible AMPylation of BiP, an Hsp70 chaperone and master regulator of the three branches of the UPR (28). During low ER stress, FicD AMPylates BiP, thereby inactivating the chaperone in the ER lumen (27, 29–31). During increased ER stress, FicD’s activity switches to BiP deAMPylation, thereby acutely increasing the pool of active BiP chaperone (32).

Previously, we established a role for FicD AMPylation of BiP in regulating the UPR of the mammalian exocrine pancreas (33). Mice lacking FicD (*FicD*^−/−^) exhibit elevated serum markers for pancreatic dysfunction, enhanced UPR signaling during fast-feeding stress, and increased fibrosis upon recovery of pharmacologically induced acute pancreatitis (33). We propose FicD-mediated AMPylation of BiP acts as a molecular rheostat for the UPR, and disruption of this regulatory mechanism upsets protein homeostasis, ultimately leading to maladaptive tissue response and increased progression of disease.

In this study, we have continued our exploration of FicD’s role in regulating UPR signaling during physiological and pathological ER stresses by focusing on the liver, a highly metabolic tissue with strong effects of UPR signaling on its physiological function (34–37). Using a transgenic reporter for UPR signaling as well as qPCR and western blot analysis, we found that livers lacking FicD exhibit enhanced UPR signaling in response to short term physiologic fasting and feeding stress. However, the livers showed no FicD-dependent changes to chronic HFD feeding or acute pathological UPR induction. Intriguingly, further analysis of repetitive pathological UPR induction suggests a role for FicD in regulating UPR responses and maintaining resiliency of tissue through adaptation to repeated ER stress. These and our previous findings are consistent with FicD being required to maintain tissue resilience during repeated physiological and pathological stresses.

## Results

### Loss of FicD results in altered UPR signaling in fasting and feeding liver

Previously, we found that *FicD*^−/−^ mice expressed elevated levels of UPR-induced transcripts in the pancreas after overnight fasting followed by 2 hours of feeding (33). Because the liver plays a critical role in regulation of metabolism both in the fasted and postprandial state, we predicted that *FicD*^−*/*−^ mice would show changes in UPR activation in this tissue under these conditions as well.

Because the liver is metabolically active during both fasting and feeding (38, 39), we first asked if the UPR was altered during the fasted state of *FicD*^−*/*−^ mice. We compared *FicD*^*fl/fl*^ and *FicD*^−*/*−^ mice that were fasted for 18 hours to *FicD*^*fl/fl*^ controls that were fed without restriction (**Fig. 1**). To assess the activation of the UPR in the liver, we examined levels of UPR-induced transcripts. Activation of ATF6 induces the expression of BiP (40). PERK activation leads to increases in CHOP and ATF4 expression (41). IRE1 catalyzes the splicing of X-box-binding protein 1(XBP1) mRNA, producing *Xbp1s* (42, 43). Levels of the UPR induced genes *BiP, Chop*, and *A14* were unchanged in both *FicD*^*fl/fl*^ and *FicD*^−*/*−^ mice during fasting (**Fig. 1A-C**). As previously reported (44), levels of *Xbp1s* were reduced in fasting livers of *FicD*^*fl/fl*^ and *FicD*^−*/*−^ mice (**Fig. 1D**). Of note, expression levels of *Xbp1s* were also reduced in *FicD*^−*/*−^ mice under control conditions, suggesting a reduction in baseline spliced Xbp1 (**Fig. 1D**). Expression of *Ficd* transcript, a UPR regulated gene, was also reduced in fasted livers of *FicD*^*fl/fl*^ mice (**Fig. 1E**).

**Figure 1.**
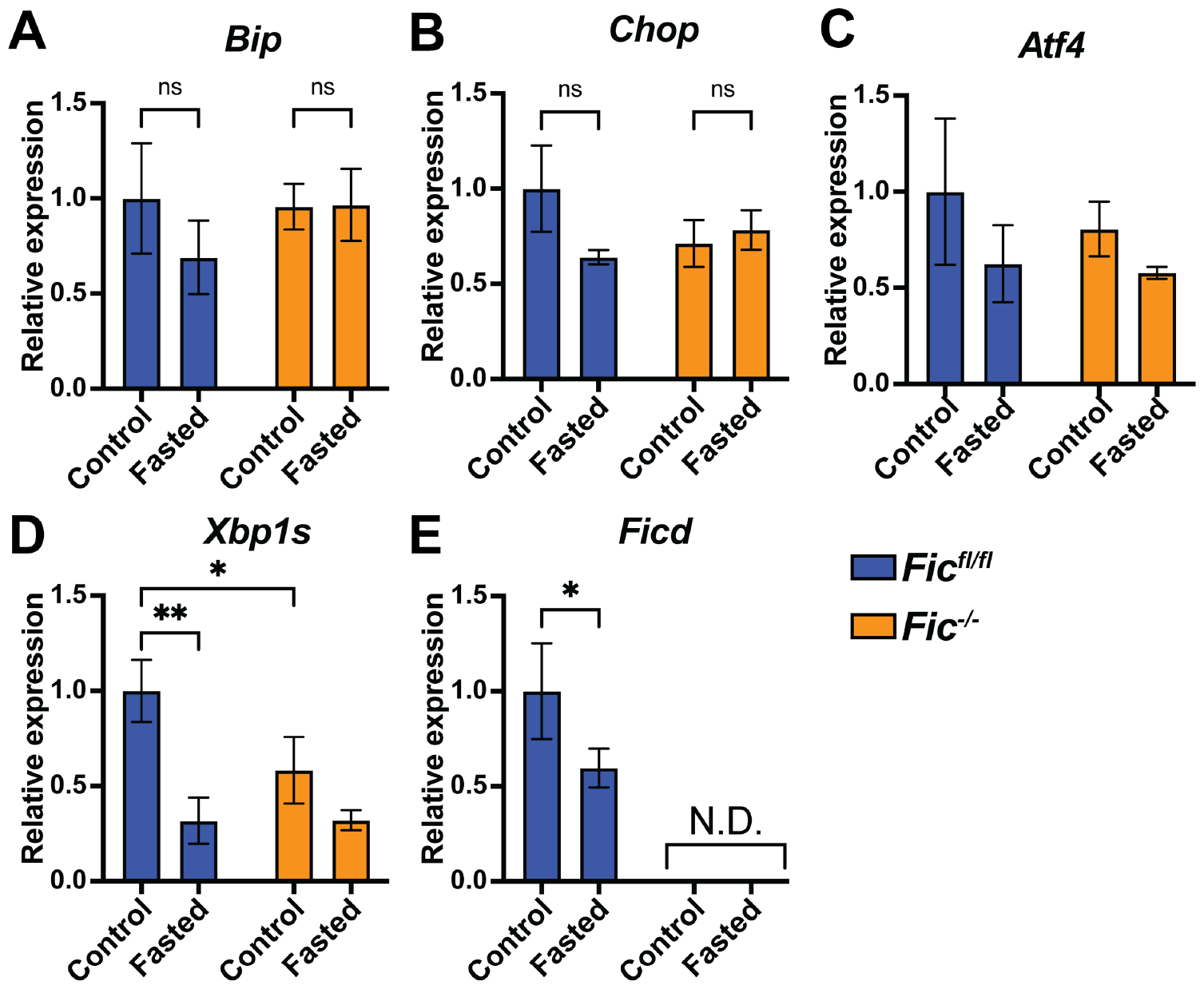
UPR signaling in fasted Fic^fl/fl^ and Fic^−/−^ liver. A-E) Quantification of *Bip, Chop, Xbp1s, A14*, and *Ficd* mRNA analyzed by qPCR from *FicD*^*fl/fl*^ (blue bar) and *FicD*^−/−^ (orange bar) mouse liver after ad libitum feeding (control) and 18hr fasting. Expression values were normalized to that of the housekeeping gene *Gapdh*. Bars indicate mean relative expression compared to fasted controls, and error bars represent standard error. Statistics were performed using GraphPad Prism 9 using an 2-way ANOVA. N=8. *, p < 0.05; **, p < 0.01. N.D., not detected

To assess the activation of the UPR in the liver during feeding, we examined levels of UPR-induced transcripts using qPCR during fasted, fasted-fed, and fasted-fed-recovery conditions as previously described (33). Consistent with previous reports of UPR induction in fasted-fed mice (45), feeding after 16-hour fasting resulted in a significant increase in UPR-regulated transcripts in the liver (**Fig. 2**). Levels of *Bip, Chop, Xbp1s*, and *A13* transcripts increased during fasted-fed conditions in both *FicD*^*fl/fl*^ and *FicD*^−*/*−^ liver (**Fig. 2A-D**). However, expression of *Chop, Xbp1s*, and *A13* transcripts were significantly increased in livers of *FicD*^−*/*−^mice compared to fasted-fed *FicD*^*fl/fl*^ controls. Levels of *A14* transcript did not increase in livers of fasted-fed *FicD*^*fl/fl*^ mice, but expression levels of *A14* transcript trended higher in livers of fasted-fed *FicD*^−*/*−^mice. Levels of *Ficd* transcript were also found to increase in the liver of *FicD*^*fl/fl*^ mice during fasting-fed conditions. Interestingly, the expression pattern of *Ficd* transcript levels remained elevated during fasted-fed-recovery in *FicD*^*fl/fl*^ livers (**Fig 2F**), like that of the *Bip* transcript expression pattern. Taken together these results suggest that *Fic*^−*/*−^ mice may be more responsive to ER stress induction during feeding after fasting.

**Figure 2.**
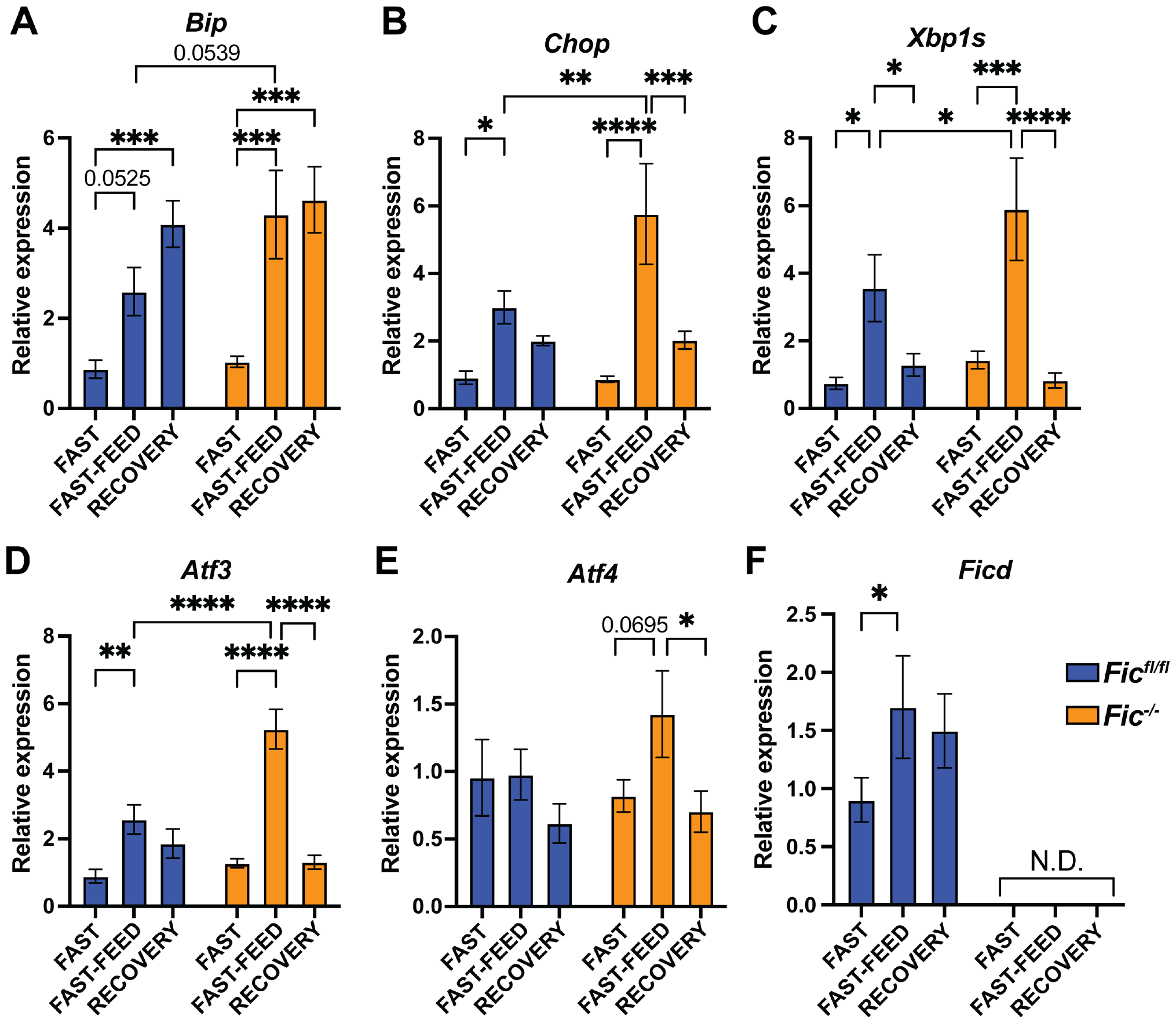
UPR signaling in fasted-fed Fic^fl/fl^ and Fic^−/−^ liver. A-F) Quantification of *Bip, Chop, Xbp1s, A13, A14*, and *Ficd* mRNA analyzed by qPCR from *FicD*^*fl/fl*^ (blue bar) and *FicD*^−/−^ (orange bar) mouse liver after fasting, fast-feeding, and fast-feed-recovery. Expression values were normalized to that of the housekeeping gene *Gapdh*. Bars indicate mean relative expression compared to fasted controls, and error bars represent standard error. Statistics were performed using GraphPad Prism 9 using an 2-way ANOVA. N=8. *, p < 0.05; **, p < 0.01, ***, p < 0.001, ****, p < 0.0001. N.D., not detected.

### Loss of FicD altered metabolism regulation in fasting and feeding liver

Next, we examined the expression patterns of UPR associated transcripts involved in regulation of metabolism in the liver, comparing changes between both control and fasting conditions as well as fasted, fasted-fed, and fasted-fed-recovery conditions.

GalE (UDP-galactose 4’ epimerase) is an enzyme responsible for the reversible conversion of UDP-galactose to UDP-glucose. *GalE* expression is linked to *Xbp1s* activation, and overexpression of GalE is linked to impaired glucose tolerance in mice (45). Consistent with previous reports, levels of *Gale* transcript were found to decrease during fasting conditions and increase during fasted-fed conditions in both *FicD*^*fl/fl*^ and *FicD*^−*/*−^ liver (**Fig. 3A & B**). However, the expression *of Gale* transcript was significantly increased in *FicD*^−*/*−^ mice after 2 hours of feeding compared to *FicD*^*fl/fl*^. This expression pattern is similar to that observed with *BiP* expression in the fasted-fed liver (Figure 2A).

**Figure 3.**
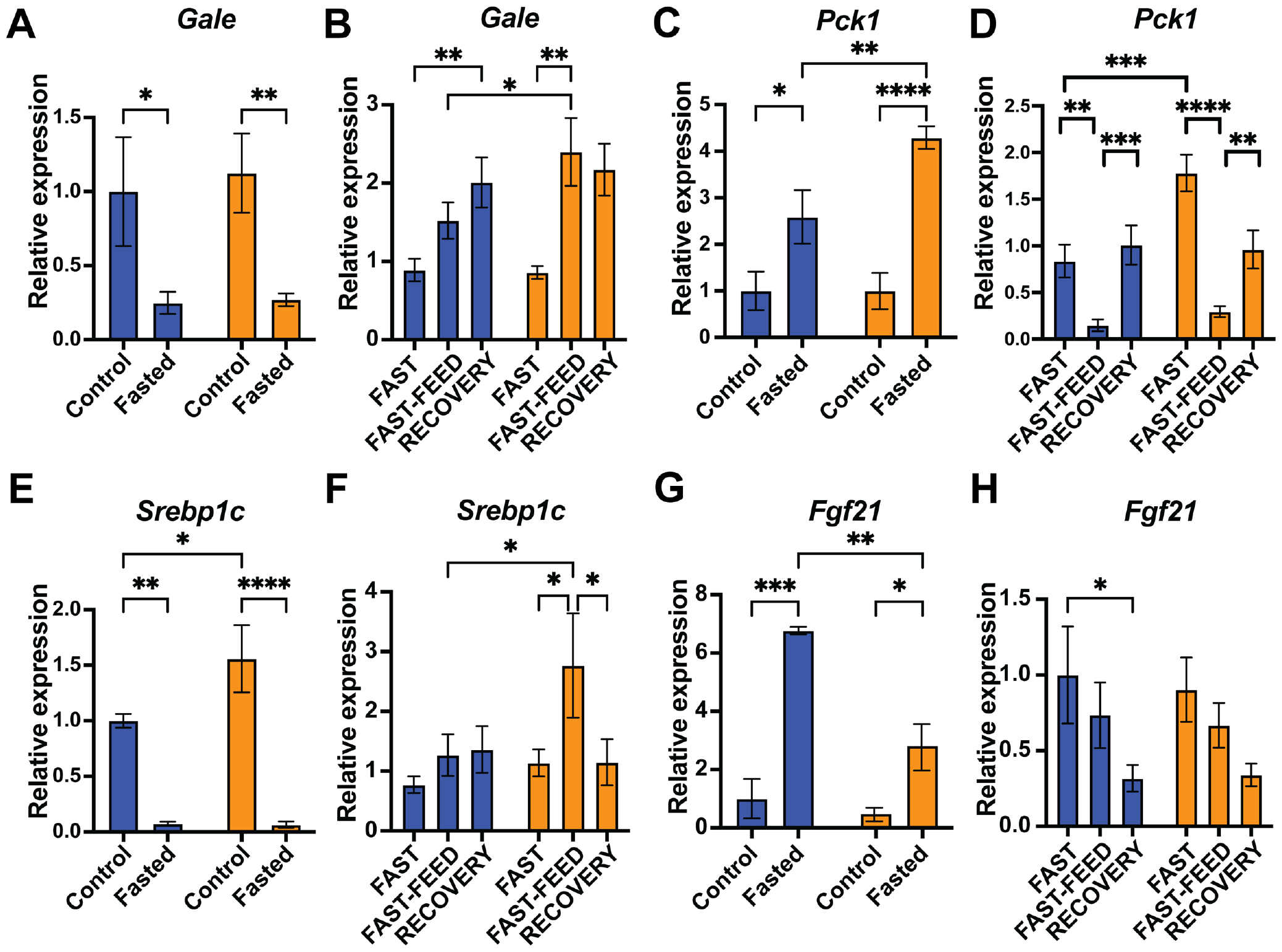
Hepatic metabolism regulation in fasted and fasted-fed Fic^fl/fl^ and Fic^−/−^ liver. A-H) Quantification of *Gale, Pck1, Srebp1c*, and *Fgf21* mRNA analyzed by qPCR from *FicD*^*fl/fl*^ (blue bar) and *FicD*^−/−^ (orange bar) mouse liver (A,C,E,G) after ad libitum feeding (control) and 18hr fasting and (B,D,F,H) fasting, fast-feeding, and fast-feed-recovery. Expression values were normalized to that of the housekeeping gene *Gapdh*. Bars indicate mean relative expression compared to fasted controls, and error bars represent standard error. Statistics were performed using GraphPad Prism 9 using an 2-way ANOVA. N=6. *, p < 0.05; **, p < 0.01, ***, p < 0.001, ****, p < 0.0001.

Pck1 (Phosphoenolpyruvate carboxykinase 1) is an enzyme involved in the regulation of gluconeogenesis. Activation of the IRE1/Xbp1s pathway of the UPR represses *Pck1* transcript expression (46). Consistent with previous studies examining Pck1 (47), levels of *Pck1* transcript increased during fasting and decreased during fasted-fed conditions in both *FicD*^*fl/fl*^ and *FicD*^−*/*−^ liver (**Fig. 3C & D**). Moreover, expression of *Pck1* transcript was significantly elevated in fasted *FicD*^−*/*−^ liver compared to fasted *FicD*^*fl/fl*^ *controls*.

SREBP-1 (sterol regulatory element-binding protein 1) is an ER regulated transcription factor with critical roles in regulating expression of metabolic genes. SREBP-1C is the predominant isoform of SREBP-1 in the liver (48). Previous reports found that *Srebpc1* transcript is downregulated during fasting and upregulated during prolonged periods of feeding (49), and UPR signaling resulted in upregulation of SREBP-1C translation via PERK pathway activation (50). Relative expression levels of *Srebp1c* transcript were found to decrease during fasting conditions and in both *FicD*^*fl/fl*^ and *FicD*^−*/*−^ liver (**Fig. 3E)**. However, *Srebp1c* transcript levels were significantly increased in the livers of *FicD*^−*/*−^ control mice that were fed without restriction compared to *FicD*^*fl/fl*^. Furthermore, 2 hours of feeding after fasting resulted in increased *Srebp1c* transcript in the livers of *FicD*^−*/*−^ but not *FicD*^*fl/fl*^ mice.

Fgf21 (Fibroblast growth factor 21) is secreted by the liver during fasting and is regulated by the IRE1α/Xbp1 branch of the UPR (51). Expression of *Fgf21* transcript was significantly reduced in the fasted livers of *FicD*^−*/*−^ mice compared to *FicD*^*fl/fl*^ *controls* (**Fig. 3G**). Levels of *Fgf21* transcript were downregulated after 2 hours of feeding in both *FicD*^*fl/fl*^ and *FicD*^−*/*−^ liver (**Fig. 3H**).

All four UPR responsive metabolic genes tested displayed expression patterns consistent with the observed changes in UPR induction in both *FicD*^*fl/fl*^ and *FicD*^−*/*−^ liver under fasting and fasting-feeding conditions. However, changes in metabolic gene expression were overall amplified in *FicD*^−*/*−^ liver. These results suggest that *FicD*^−*/*−^ liver is more responsive to metabolic changes during fasting and feeding cycles.

### UPR stress caused by prolonged high fat diet feeding is not significantly altered in FicD^−/−^ mice

Because we observed changes both in UPR and metabolic gene expression to fasting/feeding in the *FicD*^−*/*−^ liver, we next asked if the prolonged metabolic stress of long-term high fat diet (HFD) feeding would also induce significant changes to global UPR induction or metabolic regulation. To study this we carried out a longitudinal study for a high fat diet over 6 months.

Previously, we noticed that downstream transcripts regulated by the PERK branch of the UPR were particularly enhanced in *FicD*^−*/*−^ tissues and cells (22, 33, 52). PERK activation leads to global changes to translation, including enhanced translation of stress responsive genes such as the transcription factor ATF4(53–55). Therefore, we wished to monitor how loss of FicD is globally affecting PERK-induced translation of ATF4. We utilized an established reporter transgene of the PERK branch of the UPR, the UMAI luciferase reporter transgene (C57BL/6J-CAG-hATF4(uORFs)-LUC-F) (56). This UMAI reporter gene monitors the PERK branch of the UPR by indicating ATF4 translation with bioluminescence. In the UMAI reporter, under non-stress conditions translation of luciferase is predominately repressed whereas during stress conditions, the luciferase reporter is translated.

*UMAI FicD*^*+/+*^ and *UMAI FicD*^−/−^ mice were given a control diet (CD) (standard 10kcal% fat) diet or a high-fat diet (HFD) (60 kcal% fat). Animals underwent whole body imaging analysis to detect the bioluminescent UMAI-luciferase signals 0 days, 1 month, 3 months, and 6 months after the start of diet challenge (**Fig. 4A & B**). Weight gain of *UMAI FicD*^*+/+*^ and *UMAI FicD*^−/−^ was measured over 26 weeks and indicated no significant differences between *FicD*^−/−^ and *FicD*^*+/+*^ mice (**Fig. S1A)**. Whole body imaging of *UMAI FicD*^*+/+*^ and *UMAI FicD*^−/−^ demonstrated that prolonged HFD feeding results in global elevation of ATF4 translation compared to CD feeding after 3 months (**Fig. 4A & B and Fig. 4S1A**). No difference in UMAI-luciferase activity was seen between *UMAI FicD*^*+/+*^ and *UMAI FicD*^−/−^ mice on control diets during this period.

**Figure 4.**
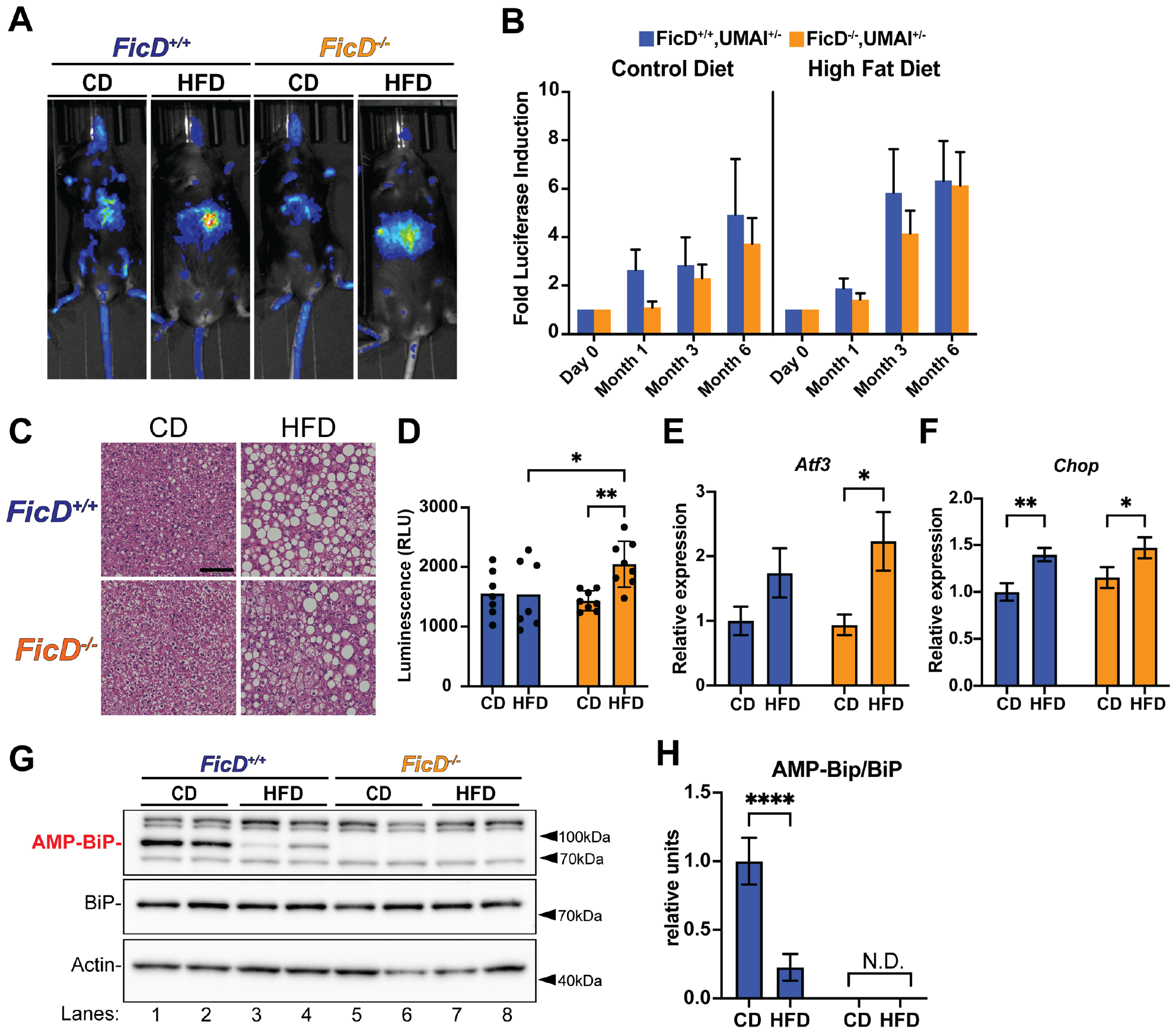
ATF4 activation during HFD feeding in FicD^fl/fl^ and FicD^−/−^ livers. A) Representative luciferase image of *FicD*^*+/+*^ *UMAI*^*+/-*^ (blue) and *FicD*^−/−^ *UMAI*^*+/-*^ mice fed CD and HFD over 6 months. B) Fold induction of luciferase activity of *FicD*^*+/+*^ *UMAI*^*+/-*^ (blue) and *FicD*^−/−^ *UMAI*^*+/-*^ mice fed CD and HFD over 6 months. N=9-13. C) Representative images of H&E stained liver of *FicD*^*+/+*^ *UMAI*^*+/-*^ and *FicD*^−/−^ *UMAI*^*+/-*^ mice fed CD and HFD over 6 months. D) Luciferase activity measured in relative luminescence units (RLU) of lysates from *Fic*^*+/+*^ *UMAI*^*+/-*^ (blue) and *FicD*^−/−^ *UMAI*^*+/-*^ (orange) liver after 6 months of control diet (CD) and high fat diet (HFD) feeding. E-F) Quantification of *A13* and *Chop* mRNA analyzed by qPCR from *FicD*^*+/+*^ *UMAI*^*+/-*^ (blue) and *FicD*^−/−^ *UMAI*^*+/-*^ (orange) mouse liver after6 months of CD and HFD feeding Expression values were normalized to that of the housekeeping gene *U36B4*. N=4-6. G) Representative western blot of protein lysates from isolated from *FicD*^*fl/fl*^ and *FicD*^−/−^ livers. Blots were probed with anti-AMP, anti-BiP, and anti-Acton antibodies. H) Quantification of relative anti-AMP to anti-BiP. N=6.Bars indicate mean relative expression compared to *Fic*^*+/+*^ *UMAI*^*+/-*^ CD controls, and error bars represent standard error. Statistics were performed using GraphPad Prism 9 using an 2-way ANOVA. *, p < 0.05; **, p < 0.01, ***, p < 0.001, ****, p < 0.0001. N.D., not detected.

To examine UPR induction in the liver, we collected liver tissue from *UMAI FicD*^*+/+*^ and *UMAI FicD*^−/−^ mice after 6 months on CD and HFD. Both *UMAI FicD*^*+/+*^ and *UMAI FicD*^−/−^ displayed increased hepatic steatosis upon HFD feeding (**Fig. 4C**). Liver tissues lysates were also assayed for the *UMAI* luciferase activity using a Firefly Luciferase Glow Assay Kit. Compared to *UMAI FicD*^*+/+*^, *UMAI FicD*^−/−^ liver lysates displayed enhanced luminescence, indicating elevated ATF4 translation in *FicD*^−/−^ liver under HFD (**Fig. 4D)**. Expression of the Atf4 target *Atf3*, but not *Chop*, was also slightly enhanced in *UMAI FicD*^−/−^ liver by qPCR (**Fig. 4D & E)**. Furthermore, expression levels of *Bip, Gale, and Srebp1c* transcripts were not significantly altered in the *FicD*^−/−^ liver (**Fig. S1B-D**). Using an anti-AMP antibody, we assessed levels of BiP AMPylation in liver lysates from HFD and CD fed mice and observed reduced BiP AMPylation in *FicD*^*+/+*^ liver on HFD compared to CD (**Fig. 4F-G**). As expected, no AMPylation or expression of FicD was observed in *FicD*^−/−^ liver (**Fig. 4H**).

### Acute UPR induction by tunicamycin is not altered in FicD^−/−^ liver

Next, we sought to assess acute UPR induction in *FicD*^−/−^ liver in response to a maladaptive, pathological ER stress. We injected mice with a single dose of tunicamycin (Tm), a well-established inducer of ER stress in the liver. In this study, 8–10-week-old male mice were injected with either a moderate dose (0.25mg/kg) of Tm, a high dose (1.0mg/kg) of Tm, or vehicle alone. Animals were sacrificed at 4hr and 24hr after administration of the drug.

Immediately upon sacrifice, serum was collected, and liver was isolated from each mouse for RNA quantification and histopathology. As previously described (34), treatment with Tm robustly induces hepatic steatosis and UPR signaling in the liver (**Fig. 5**). Both *FicD*^*fl/fl*^ and *FicD*^−/−^ livers displayed similar degrees of hepatic steatosis by Oil Red O staining (**Fig. 5A & B)** and serum analysis **(Fig. 5C and Fig. S2A-C)**.

**Figure 5.**
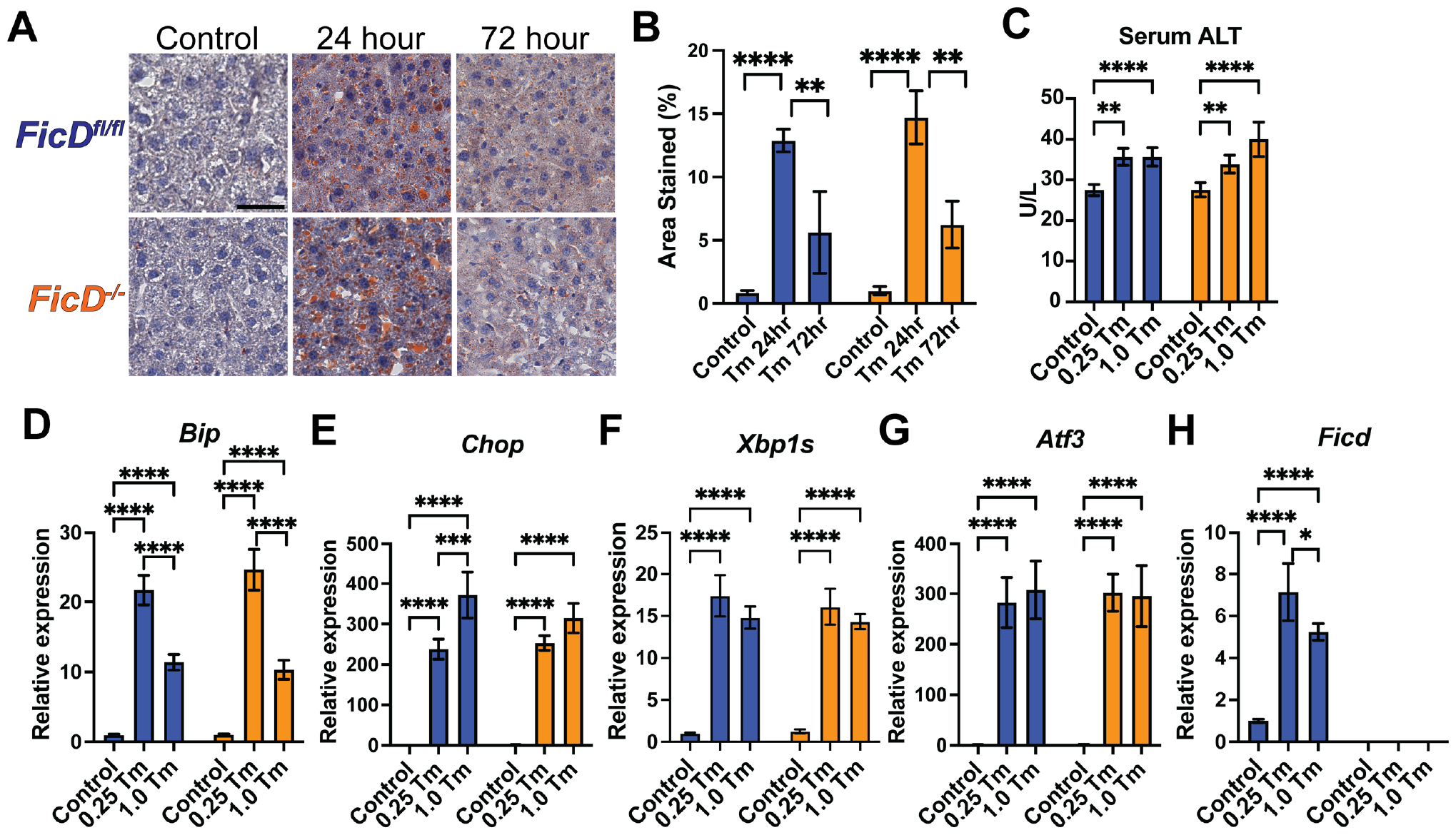
Acute UPR signaling in tunicamycin treated FicD^fl/fl^ and FicD^−/−^ liver. A) Representative images of Oil Red O stained liver of *FicD*^*fl/fl*^ and *FicD*^−/−^ mice 24 and 72 hours after treatment with 1mg/kg Tm or 24 hours vehicle control. B) Percentage of Oil Red O positive area of liver 24 and 72 hours treatment with 1mg/kg Tm and 24 hours treatment with vehicle (control). Scale bar, 50uM. C) Serum ALT analysis of *FicD*^*fl/fl*^ and *FicD*^−/−^ mice 24hours after treatment with vehicle (control), 0.25mg/kg Tunicamycin, or 1mg/kg Tm. D-H) Quantification of *Bip, Chop, Xbp1s, A13*, and *Ficd* mRNA analyzed by qPCR from *FicD*^*fl/fl*^ (blue bar) and *FicD*^−/−^ (orange bar) mouse liver 4 hours treatment with vehicle (control), 0.25mg/kg Tunicamycin, or 1mg/kg Tm. Expression values were normalized to that of the housekeeping gene *Gapdh*. Bars indicate mean relative expression compared to wildtype controls, and error bars represent standard error. Statistics were performed using GraphPad Prism 9 using an 2-way ANOVA. N=5-6. *, p < 0.05; **, p < 0.01, ***, p < 0.001, ****, p < 0.0001.

Both *FicD*^*fl/fl*^ and *FicD*^−/−^ livers displayed robust induction of UPR gene expression (*Bip, Chop, Xbp1s, A13*) at 4hrs after administration of either moderate (0.25mg/kg) or high (1.0mg/kg) doses of Tm (**Fig. 5D-G**). In *FicD*^*fl/fl*^ livers, *Ficd* transcript levels are also robustly elevated with administration of Tm (**Fig. 5H)**. Interestingly, *Bip* and *Ficd* expression was higher at 4 hours in the moderate dose of Tm as compared to expression in the high dose of Tm (**Fig. 5E & H)**.

*Gale* expression was induced in *FicD*^*fl/fl*^ and *FicD*^−/−^ livers with Tm to similar extent (**Fig. S2D**). Both *FicD*^*fl/fl*^ and *FicD*^−/−^ livers displayed similar trends in a dose dependent reduction of *Srebp1c* transcript levels with Tm treatment (**Figure S2D**). However, only *FicD*^−/−^ livers displayed statistically significant reductions in of *Srebp1c* transcript with Tm treatment, likely due to its elevated levels in untreated controls. This is consistent with recent work from the Hellerstein lab that report lipogenesis and cholesterol synthesis is repressed during UPR stress (57).

### Loss of FicD alters adaptation to repetitive acute ER stress

To determine if FicD plays a role in modulation of UPR signaling during repetitive ER stress, we assessed the luciferase production in 8–10-week-old male *UMAI FicD*^*+/+*^ and *UMAI FicD*^−/−^ mice during repetitive weekly injection of moderate dose (0.25mg/kg) Tm. Using whole body bioluminescence imaging (BLI), expression of the ATF4-Luciferase reporter was measured before injection (T=0), four hours after injection (T=4hr), and 24 hours after injection (T=24hr). This process was repeated for three weeks (**Fig. 6**). After the third and final 24-hour imaging session, mice were sacrificed, and tissues were collected from each mouse for analysis.

**Figure 6.**
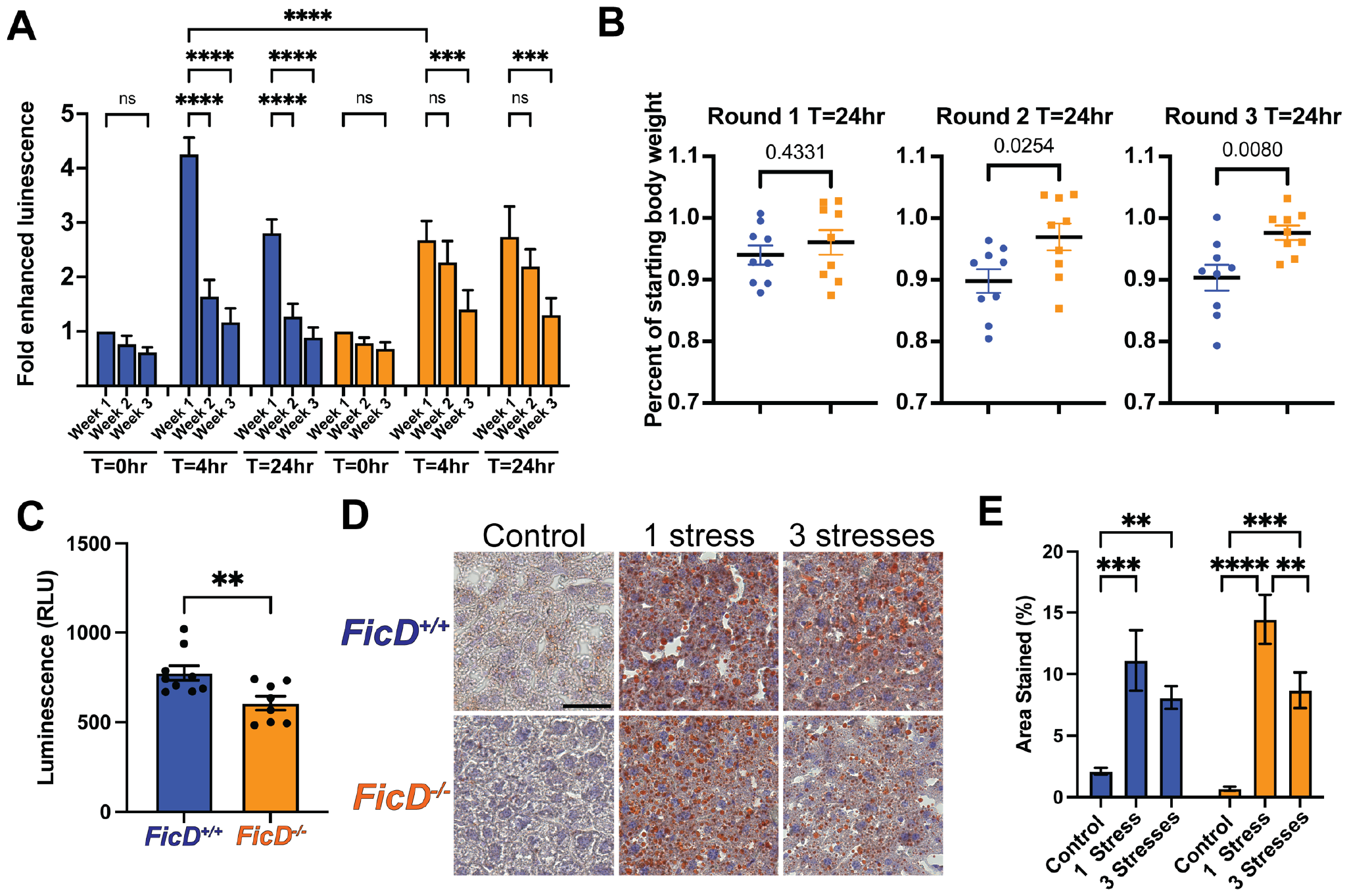
Adaptation of FicD^fl/fl^ and FicD^−/−^ mice and livers during repetitive tunicamycin challenge. A) Fold induction of total body luciferase activity of *Fic*^*+/+*^ *UMAI*^*+/-*^ (blue) and *FicD*^−/−^ *UMAI*^*+/-*^ during three weekly rounds of 0.25mg/kg tunicamycin treatment. Measurements taken at T=0, T=4, and T=24hr after injection. B) Normalized body weight of *FicD*^*+/+*^ *UMAI*^*+/-*^ (blue) and *FicD*^−/−^ *UMAI*^*+/-*^ (orange) during three weekly rounds of 0.25mg/kg tunicamycin treatment. Measurements taken at T=0, T=24, T=48hr, and T=72hr after injections. Symbols indicate mean change in body weight compared to Week 1 T=0hr timepoint. C) Luciferase activity measured in relative luminescence units (RLU) of lysates from *Fic*^*+/+*^ *UMAI*^*+/-*^ (blue) and *FicD*^−/−^ *UMAI*^*+/-*^ (orange) liver 24 hours after third dose of 0.25mg/kg tunicamycin. D) Representative images of Oil Red O stained liver of *FicD*^*fl/fl*^ and *FicD*^−/−^ mice 24 hours after one or three treatments with 0.25mg/kg Tm or vehicle control. E) Percentage of Oil Red O positive area of liver after one or three treatments with 0.25mg/kg Tm or vehicle control. Scale bar, 50uM. Statistics were performed using GraphPad Prism 9 using an student’s t test. N=8-9. ns, not significant; *, p < 0.05; **, p < 0.01, ***, p < 0.001, ****, p < 0.0001.

In *UMAI FicD*^*+/+*^ *mice*, we observed a strong initial induction of ATF4-Luciferase reporter at the T=4hr and T=24hr during the first (week 1) round of treatment. However, when the Tm treatment was repeated, *UMAI FicD*^*+/+*^ mice displayed a marked reduction in ATF4-Luciferase expression **(Fig. 6A and Fig. S3A)**. Conversely, in *UMAI FicD*^−/−^ mice we observed a weaker initial induction of ATF4-Luciferase during the first round of treatment. When the Tm treatment was repeated a second time in *UMAI FicD*^−/−^ mice, we did not observe a significant decrease in ATF4-Luciferase expression at the T=4hr and T=24hr. However, a decrease in ATF4-Luciferase induction was observed in *UMAI FicD*^−/−^ mice during the third round of Tm treatment.

We also observed a striking difference in the patterns of weight loss between *UMAI FicD*^*+/+*^ and *UMAI FicD*^−/−^ mice during repetitive Tm stress. Both *UMAI FicD*^*+/+*^ and *UMAIFicD*^−/−^mice steadily lost body weight for the first 72 hours during the initial Tm stress treatment **(Fig. 6B and Fig. S3B)**, eventually regaining all lost body weight by one week of recovery. During the second and third Tm stress event, *UMAI FicD*^*+/+*^ control mice experienced a very sharp decline in body weight loss within 24 hours of Tm challenge. However, *UMAI FicD*^−/−^ mice experienced a slower, steadier decline in weight, more consistent with weight loss patterns observed during the initial Tm stress. Taken together, this data suggests that *FicD*^−/−^ mice may lack regulatory mechanisms for tempering or adapting to repetitive challenges to ER homeostasis and cellular health.

### Loss of FicD does not contribute to liver damage with repetitive Tm challenge

To look more closely at the effect of repetitive Tm treatment on the liver, we assayed liver lysates for the *UMAI* luciferase activity using a Firefly Luciferase Glow Assay Kit. Despite the increase in total body ATF4-Luciferase observed for *UMAI FicD*^−/−^ mice after three challenges with Tm, liver lysates from *UMAI FicD*^−/−^ mice displayed reduced luminescence compared to *UMAI FicD*^*+/+*^ controls 24 hours after the repetitive Tm challenges. This data suggests that *FicD*^−/−^ liver may have reduced PERK branch UPR signaling under repetitive stress (**Fig. 6C)**.

To determine if loss of FicD resulted in altered effects on the liver tissue during repetitive ER stress, liver sections from the repetitively stressed mice were analyzed for histopathological changes. As previously observed, 24hrs after both a single and repetitive Tm stress, we see increased micro steatosis in both *UMAI FicD*^*+/+*^ and *UMAI FicD*^−/−^ livers. Furthermore, both *UMAI FicD*^*+/+*^ and *UMAI FicD*^−/−^ livers exhibited areas of mild centrilobular necrosis (death of hepatocytes around the central veins); however, no appreciable difference in necrosis or hepatic steatosis was observed between genotypes at either timepoint (**Fig. 6D & E**).

## Discussion

Previously, we showed that loss of FicD regulation in the pancreas led to changes in UPR signaling during fasting and feeding (33). The loss of FicD caused damage to the pancreas when animals were treated with a repeated stress. This observation was also seen Drosophila eyes that lacked FicD and were treated with repeated light stress (22). Although, we suspected other tissues with UPR regulated responses might have similar effects to this physiological stimulus, we wanted to test an organ with regenerative capacity. The liver, unlike the pancreas, is endowed with this ability and therefore was an intuitive next choice to study.

We found that UPR signaling during fasting/feeding was indeed enhanced in *FicD*^−*/*−^ livers. Furthermore, we found *FicD*^−*/*−^ livers exhibited amplified transcriptional responses in metabolic genes involved in glucose and lipid metabolism. Of note, these patterns of expression remained largely consistent with wildtype liver, but, like UPR signaling, the sensitivity and magnitude of the responses were enhanced in *FicD*^−/−^ liver. These data indicate that FicD plays a role in tempering UPR signaling during mild, transitory physiological stresses (**Fig. 2**)

Due to the link between enhanced UPR signaling and changes in liver metabolism, we wanted to assess the role of FicD in the regulation of prolonged metabolic stress in the liver. To do this, we monitored expression of an ATF4 reporter transgene (UMAI) in *FicD*^−/−^ and wildtype control mice during 6 months of high fat feeding. Globally, we found no significant change in ATF4 translation during 6 months of high fat feeding. However, our analysis revealed enhanced ATF4 translation and UPR signaling in *FicD*^−/−^ livers on HFD with no change in metabolic regulation between *FicD*^*fl/fl*^ and *FicD*^−/−^ (**Fig. 4**).

Next, we wanted to determine if FicD played a role in the regulation of stronger, more pathological ER stressors. To do this, we assessed the effects of a single moderate or high dose of Tm on the liver. Our transcriptional analysis indicated that both *FicD*^*fl/fl*^ and *FicD*^−*/*−^ livers elicit robust and equivalent acute UPR responses to Tm with equivalent pathological stress occurring in *FicD*^*fl/fl*^ and *FicD*^−*/*−^ livers. (**Fig. 5**)

Finally, we wanted to determine if FicD played a role in protecting the liver from repetitive acute ER stress. To do this, we used repetitive Tm and recovery regimen in wildtype *UMAI FicD*^*+/+*^ and *UMAI FicD*^−/−^ ATF4 reporter mice. Consistent with previous reports using very low dose chronic Tm inductions (58), we found that wildtype mice displayed a progressive reduction in UPR signaling with repetitive treatment of moderate Tm, indicative of adaptation to a repetitive ER stress. However, *FicD*^−/−^mice failed to exhibit this adaption to Tm stress, as indicated by elevated total body ATF4-Luciferease activity with repetitive stress.

Intriguingly, loss of this stress adaptation appeared to some extent beneficial to *FicD*^−/−^ mice. Wildtype mice exhibited increased rapid weight loss within 24 hours of the second and third Tm treatments, while *FicD*^−/−^ lost weight more slowly and to a lesser degree. Of note, the level of ATF4-Luciferease activity observed in full body imaging of *FicD*^−/−^ mice under chronic and repetitive stresses was elevated compared to wildtype controls; however, measured levels of ATF4-Luciferease activity were lower in *FicD*^−/−^ liver. Histology of liver showed no significant change in liver pathology between *UMAI FicD*^*+/+*^ and *UMAI FicD*^−/−^.

It’s interesting to note that expression of the *Srebp1c* transcript was elevated in *FicD*^−/−^ mice under control conditions (Figure 3E, Figure S5E) and was found to be more responsive to feeding after fasting in *FicD*^−/−^ mice where UPR induction is elevated. However strong UPR induction with Tm results in decreased *Srebp1c* transcript, indicating Srebp1c may be positively regulated by mild, adaptive UPR signaling but negatively regulated by stronger UPR inductions.

This study, when combined with findings from our previous studies in exocrine pancreas, suggests that FicD regulates UPR in two ways. First, our data indicate that FicD plays a role in tempering UPR signaling during mild, transitory physiological stresses in the liver, such as in fasting/feeding. Second, our repetitive tunicamycin studies suggest a role for FicD in the recovery of and adaptation to repetitive stresses. For the latter, this discrepancy indicates that other tissues, as observed with the pancreas, could be more severely or differentially affected by the loss of FicD, potentially contributing to the observed weight loss phenotype.

Though we found some changes to UPR signaling and metabolic gene regulation, livers remained remarkably resilient in the absence of *FicD*. The liver is continuously metabolically active, so the role of FicD may be diminished in this robust tissue with little metabolic down time. We think our observed tissue specific differences are an indication that the role of FicD may be both tissue and stressor dependent and partially redundant with other key UPR regulatory mechanisms of some tissues.

Our current studies are consistent with our previous observations that the PERK branch of the UPR is linked with FicD activity *in vivo* (22, 33, 52). Because PERK activation leads to global changes to translation, tissues with high fluctuations in secretory demands may be more reliant on FicD regulation of BiP Future studies will focus on secretory tissues with established links to UPR dysregulation and disease, such as the brain. Two point mutations in FicD have been identified in the human population with connections to neuronal defects (59, 60). Determining how FicD regulation of UPR signaling affects neuronal function and resiliency is an exciting avenue for further research.

### Experimental procedures

#### Mice

The Institutional Animal Care and Use Committee of the University of Texas Southwestern Medical Center approved all experiments. FicD^fl/fl^ and FicD^−/−^ mouse lines are previously described (33).The UMAI luciferase reporter (C57BL/6J-CAG-hATF4(uORFs)-LUC-F) transgenic mouse line was generously provided by the laboratory of Esta Sterneck, with permission from Dr. Takao Iwawaki and Kanazawa Medical University. Mice were housed in a specific pathogen-free facility and fed a standard chow diet (#2016 Harlan Teklad), unless specified. Fasting and fasting feeding studies were performed as previously described (33). For high fat diet studies, animals were provided specialty a high fat diet (D12492, 60 kcal% fat, Research Diets, New Brunswick, NJ) or control diet (D12450J, 10 kcal% fat, Research Diets, New Brunswick, NJ). For Tm treatment, mice were administered a single intraperitoneal dose (0.25mg/kg or 1.0mg/kg) of Tunicamycin dissolved in 150mM dextrose. Mice in the control group were injected with vehicle alone. For serum analysis, blood was collected in Microvette serum collection tubes (Sarstedt) and centrifuged at 10,000xg at 4°C for 15 minutes. Serum analysis was performed by *UT Southwestern Metabolic Phenotyping Core* using VITROS MicroSlide™ technology.

#### In vivo and ex vivo luciferase detection

To detect the bioluminescence of UMAI-Luc transgenic mice, mice were imaged using an IVIS spectrum. Mice were anesthetized with 1-3% 1-2L/min Isoflurane via Precision Vaporizer. Luciferin (125mg/kg) was injected subcutaneously and imaged 5 and 10 minutes after luciferin administration. Luminescence images were captured and equal sized regions of interest (ROIs) were drawn over the torso of each mouse using Aura Imaging software, and photos/s were calculated. For studies in which luciferase was performed on ex vivo liver tissue, the Pierce™ Firefly Luciferase Glow Assay Kit (Thermo Fisher) was used on freshly lysed tissue. Bioluminescence was measured on normalized samples with CLARIOstar Plus fluorescence plate reader.

#### Histology

Mouse livers were harvested and fixed in 10% neutral buffered formalin overnight at 4°C. Paraffin sections were embedded by the UT Southwestern (UTSW) Molecular Pathology Core. Cryoembedded livers were equilibrated in 30% sucrose prior to freezing, sectioning, and staining. Hematoxylin and eosin (H&E) and Oil-Red O staining was performed using standard techniques.

#### Quantitative real-time PCR

RNA from liver was extracted using RNA Stat-60 (Iso-Tex Diagnostics). Complementary DNA (cDNA) was generated from RNA (2 μg) using the High-Capacity cDNA Reverse Transcription Kit (Life Technologies). qPCR was performed by the SYBR Green method (61). Primer sequences for the genes analyzed can be found in **Table S1**. Experiments were performed on a BioRad CFX Touch and analyzed with CFX Maestro Software.

#### Western Blot Analysis

Liver was washed in PBS and homogenized in RIPA buffer (50mM Tris pH 8, 150mM NaCl, 1% NP-40, 0.5% Na deoxycholate, PMSF, PhosSTOP (Roche), Protease Inhibitor Cocktail (Roche. Lysates were centrifuged at 10,000xg for 10 minutes to remove nuclei and cellular debris. Lysates were separated by SDS-PAGE and transferred to PVDF membranes. Blots were probed with anti-AMP 17g6 (gift from Aymelt Itzen), anti-GRP78 (Abcam), anti-actin (Sigma), anti-eif2A (Cell Signaling), anti-EIF2S1-phospho (Abcam), anti-IRE1 (Cell Signaling) Membranes were then incubated with horseradish peroxidase–conjugated secondary antibodies (Cell Signaling Technology) against the primary antibody’s host species for 1 hour. Membranes were developed using the ECL substrate solution (Bio-Rad). Quantification of western blots was performed using NIH ImageJ software. Band densities were measured and subtracted from background.

## Data availability

All data described herein are contained within the manuscript.

## Supporting information

This article contains supporting information.

## Acknowledgments

We want to thank UT Southwestern *Metabolic Phenotyping Core* for the analysis of serum metabolites and the UT Southwestern small animal imaging center for use of whole animal luciferase imaging instruments. We thank Dr. Takao Iwawaki and Kanazawa Medical University for allowing us the use of the UMAI transgenic mouse line and for Esta Sterneck for sending us the UMAI mice. We also thank Aymelt Itzen for his generous gift of monoclonal α-AMP antibodies. We thank the Helmut Krämer and Orth lab members for discussions.

## Funding and additional information

KO is a WW Caruth, Jr. Biomedical Scholar with an Earl A Forsythe Chair in Biomedical Science. Research reported in this study was supported by the Welch Foundation grant I-1561 (KO), Once Upon a Time…Foundation (KO), the National Institutes of Health Grant R35 GM134945 (KO), the National Institutes of Health Grant R21 1R21EY034597-01A1 (AKC) and UTSWNORC grant under NIDDK/NIH award number P30DK127984 (AKC). The funders had no role in study design, data collection and interpretation, or the decision to submit the work for publication.

## References

1. Luoma, P. V. (2013) Elimination of endoplasmic reticulum stress and cardiovascular, type 2 diabetic, and other metabolic diseases Ann Med 45, 194–202 10.3109/07853890.2012.700116

2. Carreras-Sureda, A., Pihan, P., and Hetz, C. (2018) Calcium signaling at the endoplasmic reticulum: fine-tuning stress responses Cell Calcium 70, 24–31 10.1016/j.ceca.2017.08.004

3. Bommiasamy, H., and Popko, B. (2011) Animal models in the study of the unfolded protein response Methods Enzymol 491, 91–109 10.1016/B978-0-12-385928-0.00006-7

4. Garg, A. D., Kaczmarek, A., Krysko, O., Vandenabeele, P., Krysko, D. V., and Agostinis, P. (2012) ER stress-induced inflammation: does it aid or impede disease progression? Trends Mol Med 18, 589–598 10.1016/j.molmed.2012.06.010

5. Thorp, E., Iwawaki, T., Miura, M., and Tabas, I. (2011) A reporter for tracking the UPR in vivo reveals patterns of temporal and cellular stress during atherosclerotic progression J Lipid Res 52, 1033–1038 10.1194/jlr.D012492

6. Batista, A., Rodvold, J. J., Xian, S., Searles, S. C., Lew, A., Iwawaki, T. et al. (2020) IREtialpha regulates macrophage polarization, PD-L1 expression, and tumor survival PLoS Biol 18, e3000687 10.1371/journal.pbio.3000687

7. Huang, N., Yu, Y., and Qiao, J. (2017) Dual role for the unfolded protein response in the ovary: adaption and apoptosis Protein Cell 8, 14–24 10.1007/s13238-016-0312-3

8. Akhter, M. S., Uddin, M. A., Kubra, K. T., and Barabutis, N. (2020) Autophagy, Unfolded Protein Response and Lung Disease Curr Res Cell Biol 1, 10.1016/j.crcbio.2020.100003

9. Geraghty, P., Wallace, A., and D’Armiento, J. M. (2011) Induction of the unfolded protein response by cigarette smoke is primarily an activating transcription factor 4-C/EBP homologous protein mediated process Int J Chron Obstruct Pulmon Dis 6, 309–319 10.2147/COPD.S19599

10. Lee, J., and Ozcan, U. (2014) Unfolded protein response signaling and metabolic diseases J Biol Chem 289, 1203–1211 10.1074/jbc.R113.534743

11. Gardner, B. M., Pincus, D., Gotthardt, K., Gallagher, C. M., and Walter, P. (2013) Endoplasmic Reticulum Stress Sensing in the Unfolded Protein Response Cold Spring Harbor Perspect Biol 5, a013169 10.1101/cshperspect.a013169

12. Bertolotti, A., Zhang, Y., Hendershot, L. M., Harding, H. P., and Ron, D. (2000) Dynamic interaction of BiP and ER stress transducers in the unfolded-protein response Nat Cell Biol 2, 326–332,

13. Walter, P., and Ron, D. (2011) The Unfolded Protein Response: From Stress Pathway to Homeostatic Regulation Science 334, 1081–1086 10.1126/science.1209038.

14. Meares, G. P., Hughes, K. J., Naatz, A., Papa, F. R., Urano, F., Hansen, P. A. et al. (2011) IRE1-dependent activation of AMPK in response to nitric oxide Mol Cell Biol 31, 4286–4297 10.1128/MCB.05668-11

15. Adachi, Y., Yamamoto, K., Okada, T., Yoshida, H., Harada, A., and Mori, K. (2008) ATF6 Is a Transcription Factor Specializing in the Regulation of Quality Control Proteins in the Endoplasmic Reticulum Cell Struct Funct 33, 75–89,

16. Mitra, S., and Ryoo, H. D. (2019) The unfolded protein response in metazoan development J Cell Sci 132, 10.1242/jcs.217216

17. Liu, L., Zhao, M., Jin, X., Ney, G., Yang, K. B., Peng, F. et al. (2019) Adaptive endoplasmic reticulum stress signalling via IREtialpha-XBP1 preserves self-renewal of haematopoietic and pre-leukaemic stem cells Nat Cell Biol 21, 328–337 10.1038/s41556-019-0285-6

18. Rutkowski, D. T., and Hegde, R. S. (2010) Regulation of basal cellular physiology by the homeostatic unfolded protein response J Cell Biol 189, 783–794 10.1083/jcb.201003138

19. Wu, J., Ruas, J. L., Estall, J. L., Rasbach, K. A., Choi, J. H., Ye, L. et al. (2011) The unfolded protein response mediates adaptation to exercise in skeletal muscle through a PGC-1alpha/ATF6alpha complex Cell Metab 13, 160–169 10.1016/j.cmet.2011.01.003

20. Salminen, A., and Kaarniranta, K. (2010) ER stress and hormetic regulation of the aging process Ageing Res Rev 9, 211–217 10.1016/j.arr.2010.04.003

21. Mollereau, B., Manie, S., and Napoletano, F. (2014) Getting the better of ER stress J Cell Commun Signal 8, 311–321 10.1007/s12079-014-0251-9

22. Moehlman, A. T., Casey, A. K., Servage, K., Orth, K., and Kramer, H. (2018) Adaptation to constant light requires Fic-mediated AMPylation of BiP to protect against reversible photoreceptor degeneration Elife 7, 10.7554/eLife.38752

23. Meyerovich, K., Ortis, F., Allagnat, F., and Cardozo, A. K. (2016) Endoplasmic reticulum stress and the unfolded protein response in pancreatic islet inflammation J Mol Endocrinol 57, R1–R17 10.1530/JME-15-0306

24. Harding, H. P., Zeng, H., Zhang, Y., Jungreis, R., Chung, P., Plesken, H. et al. (2001) Diabetes Mellitus and Exocrine Pancreatic Dysfunction in Perk-/-Mice Reveals a Role for Translational Control in Secretory Cell Survival Molecular Cell 7, 1153–1163,

25. Gomez, E., Powell, M. L., Bevington, A., and Herbert, T. P. (2008) A decrease in cellular energy status stimulates PERK-dependent eIF2alpha phosphorylation and regulates protein synthesis in pancreatic beta-cells Biochem J 410, 485–493 10.1042/BJ20071367

26. Dudek, J., Benedix, J., Cappel, S., Greiner, M., Jalal, C., Muller, L. et al. (2009) Functions and pathologies of BiP and its interaction partners Cell Mol Life Sci 66, 1556–1569 10.1007/s00018-009-8745-y

27. Worby, C. A., Mattoo, S., Kruger, R. P., Corbeil, L. B., Koller, A., Mendez, J. C. et al. (2009) The Fic Domain: Regulation of Cell Signaling by Adenylylation Mol Cell 34, 93–103 10.1016/j.molcel.2009.03.008

28. Ham, H., Woolery, A. R., Tracy, C., Stenesen, D., Kramer, H., and Orth, K. (2014) Unfolded Protein Response-regulated Drosophila Fic (dFic) Protein Reversibly AMPylates BiP Chaperone during Endoplasmic Reticulum Homeostasis J Biol Chem 289, 36059–36069 10.1074/jbc.M114.612515

29. Truttmann, M. C., Cruz, V. E., Guo, X., Engert, C., Schwartz, T. U., and Ploegh, H. L. (2016) The Caenorhabditis elegans Protein FIC-1 Is an AMPylase That Covalently Modifies Heat-Shock 70 Family Proteins, Translation Elongation Factors and Histones PLoS Genet 12, e1006023 10.1371/journal.pgen.1006023

30. Preissler, S., Rato, C., Chen, R., Antrobus, R., Ding, S., Fearnley, I. M. et al. (2015) AMPylation Matches BiP Activity to Client Protein Load in the Endoplasmic Reticulum Elife 4, e12621 10.7554/eLife.12621

31. Preissler, S., Rohland, L., Yan, Y., Chen, R., Read, R. J., and Ron, D. (2017) AMPylation targets the rate-limiting step of BiP’s ATPase cycle for its functional inactivation Elife 6, 10.7554/eLife.29428

32. Casey, A. K., Moehlman, A. T., Zhang, J., Servage, K. A., Kramer, H., and Orth, K. (2017) Fic-mediated deAMPylation is not dependent on homodimerization and rescues toxic AMPylation in flies J Biol Chem 292, 21193–21204 10.1074/jbc.M117.799296

33. Casey, A. K., Gray, H. F., Chimalapati, S., Hernandez, G., Moehlman, A. T., Stewart, N. et al. (2022) Fic-mediated AMPylation tempers the unfolded protein response during physiological stress Proc Natl Acad Sci U S A 119, e2208317119 10.1073/pnas.2208317119

34. Yamamoto, K., Takahara, K., Oyadomari, S., Okada, T., Sato, T., Harada, A. et al. (2010) Induction of liver steatosis and lipid droplet formation in ATF6alpha-knockout mice burdened with pharmacological endoplasmic reticulum stress Mol Biol Cell 21, 2975–2986 10.1091/mbc.E09-02-0133

35. Zhu, Y., Zhao, S., Deng, Y., Gordillo, R., Ghaben, A. L., Shao, M. et al. (2017) Hepatic GALE Regulates Whole-Body Glucose Homeostasis by Modulating Tff3 Expression Diabetes 66, 2789–2799 10.2337/db17-0323

36. Lee, A. H., Scapa, E. F., Cohen, D. E., and Glimcher, L. H. (2008) Regulation of Hepatic Lipogenesis by the Transcription Factor XBP1 Science 320, 1492–1496,

37. Zheng, Z., Zhang, C., and Zhang, K. (2010) Role of unfolded protein response in lipogenesis World J Hepatol 2, 203–207 10.4254/wjh.v2.i6.203

38. Rui, L. (2014) Energy metabolism in the liver Compr Physiol 4, 177–197 10.1002/cphy.c130024

39. Bideyan, L., Nagari, R., and Tontonoz, P. (2021) Hepatic transcriptional responses to fasting and feeding Genes Dev 35, 635–657 10.1101/gad.348340.121

40. Baumeister, P., Luo, S., Skarnes, W. C., Sui, G., Seto, E., Shi, Y. et al. (2005) Endoplasmic reticulum stress induction of the Grp78/BiP promoter: activating mechanisms mediated by YY1 and its interactive chromatin modifiers Mol Cell Biol 25, 4529–4540 10.1128/MCB.25.11.4529-4540.2005

41. Oyadomari, S., and Mori, M. (2004) Roles of CHOP/GADD153 in endoplasmic reticulum stress Cell Death Differ 11, 381–389 10.1038/sj.cdd.4401373

42. Lee, K., Tirasophon, W., Shen, X., Michalak, M., Prywes, R., Okada, T. et al. (2002) IRE1-mediated unconventional mRNA splicing and S2P-mediated ATF6 cleavage merge to regulate XBP1 in signaling the unfolded protein response Genes Dev 16, 452–466 10.1101/gad.964702

43. Calfon, M., Zeng, H., Urano, F., Till, J. H., Hubbard, S. R., Harding, H. P. et al. (2002) IRE1 couples endoplasmic reticulum load to secretory capacity by processing the XBP-1 mRNA Nature 415, 92–96 10.1038/415092a

44. Shao, M., Shan, B., Liu, Y., Deng, Y., Yan, C., Wu, Y. et al. (2014) Hepatic IRE1alpha regulates fasting-induced metabolic adaptive programs through the XBP1s-PPARalpha axis signalling Nat Commun 5, 3528 10.1038/ncomms4528

45. Deng, Y., Wang, Z. V., Tao, C., Gao, N., Holland, W. L., Ferdous, A. et al. (2013) The Xbptis/GalE axis links ER stress to postprandial hepatic metabolism J Clin Invest 123, 455–468 10.1172/JCI62819

46. Madhavan, A., Kok, B. P., Rius, B., Grandjean, J. M. D., Alabi, A., Albert, V. et al. (2022) Pharmacologic IRE1/XBP1s activation promotes systemic adaptive remodeling in obesity Nat Commun 13, 608 10.1038/s41467-022-28271-2

47. Sellers, J., Brooks, A., Fernando, S., Westenberger, G., Junkins, S., Smith, S. et al. (2021) Fasting-Induced Upregulation of MKP-1 Modulates the Hepatic Response to Feeding Nutrients 13, 10.3390/nu13113941

48. Shimomura, I., Shimano, H., Horton, J. D., Goldstein, J. L., and Brown, M. S. (1997) Differential Expression of Exons 1a and 1c in mRNAs for Sterol Regulatory Element Binding Protein-1 in Human and Mouse Organs and Cultured Cells J Clin Invest 99, 838–845,

49. Liang, G., Yang, J., Horton, J. D., Hammer, R. E., Goldstein, J. L., and Brown, M. S. (2002) Diminished hepatic response to fasting/refeeding and liver X receptor agonists in mice with selective deficiency of sterol regulatory element-binding protein-1c J Biol Chem 277, 9520–9528 10.1074/jbc.M111421200

50. Colgan, S. M., Tang, D., Werstuck, G. H., and Austin, R. C. (2007) Endoplasmic reticulum stress causes the activation of sterol regulatory element binding protein-2 Int J Biochem Cell Biol 39, 1843–1851 10.1016/j.biocel.2007.05.002

51. Jiang, S., Yan, C., Fang, Q. C., Shao, M. L., Zhang, Y. L., Liu, Y. et al. (2014) Fibroblast growth factor 21 is regulated by the IRE1alpha-XBP1 branch of the unfolded protein response and counteracts endoplasmic reticulum stress-induced hepatic steatosis J Biol Chem 289, 29751–29765 10.1074/jbc.M114.565960

52. Gulen, B., Kinch, L. N., Servage, K. A., Blevins, A., Stewart, N. M., Gray, H. F. et al. (2024) FicD Sensitizes Cellular Response to Glucose Fluctuations in Mouse Embryonic Fibroblasts bioRxiv 10.1101/2024.01.22.576705

53. Harding, H. P., Zhang, Y., Bertolotti, A., Zeng, H., and Ron, D. (2000) Perk Is Essential for Translational Regulation and Cell Survival during the Unfolded Protein Response Mol Cell 5, 897–904,

54. Vattem, K. M., and Wek, R. C. (2004) Reinitia1on involving upstream ORFs regulates ATF4 mRNA translation in mammalian cells Proc Natl Acad Sci U S A 101, 11269–11274 10.1073/pnas.0400541101

55. Harding, H. P., Zhang, Y., and Ron, D. (1999) Protein translation and folding are coupled by an endoplasmic-reticulum-resident kinase Nature 397, 271–274,

56. Iwawaki, T., Akai, R., Toyoshima, T., Takeda, N., Ishikawa, T. O., and Yamamura, K. I. (2017) Transgenic mouse model for imaging of ATF4 translational activation-related cellular stress responses in vivo Sci Rep 7, 46230 10.1038/srep46230

57. Ward, C. P., Peng, L., Yuen, S., Chang, M., Karapetyan, R., Nyangau, E. et al. (2022) ER Unfolded Protein Response in Liver In Vivo Is Characterized by Reduced, Not Increased, De Novo Lipogenesis and Cholesterol Synthesis Rates with Uptake of Fatty Acids from Adipose Tissue: Integrated Gene Expression, Translation Rates and Metabolic Fluxes Int J Mol Sci 23, 10.3390/ijms23031073

58. Gomez, J. A., and Rutkowski, D. T. (2016) Experimental reconstitution of chronic ER stress in the liver reveals feedback suppression of BiP mRNA expression Elife 5, 10.7554/eLife.20390

59. Rebelo, A. P., Ruiz, A., Dohrn, M. F., Wayand, M., Farooq, A., Danzi, M. C. et al. (2022) BiP inactivation due to loss of the deAMPylation function of FICD causes a motor neuron disease Genet Med 10.1016/j.gim.2022.08.019

60. Perera, L. A., Hattersley, A. T., Harding, H. P., Wakeling, M. N., Flanagan, S. E., Mohsina, I. et al. (2023) Infancy-onset diabetes caused by de-regulated AMPylation of the human endoplasmic reticulum chaperone BiP EMBO Mol Med 15, e16491 10.15252/emmm.202216491

61. Bookout, A. L., Cummins, C. L., Mangelsdorf, D. J., Pesola, J. M., and Kramer, M. F. (2006) High-Throughput Real-Time QuantitativeReverse Transcription PCR Curr Protoc Mol Bio 73,

