## Supplementary material for "FicD regulates adaptation to the unfolded protein response in the murine liver": Supp data

Affiliations
[1]Department of Molecular Biology, University of Texas Southwestern Medical Center, Dallas, TX, 75390, USA
[2]Howard Hughes Medical Institute, Dallas, TX, 75390, USA
[3]Washington and Lee University, Lexington VA 24450, USA
[4]Department of Biochemistry, University of Texas Southwestern Medical Center, Dallas, TX, 75390, USA

**Figure S1.** ***Gene expression in HFD feeding in FicD$^{fl/fl}$ and FicD$^{-/-}$ livers.***
**Figure S2.** ***Hepatic metabolism regulation in tunicamycin treated FicD$^{fl/fl}$ and FicD$^{-/-}$ liver.***
**Figure S3. Changes** ***during repetitive tunicamycin challenge and recovery in FicD$^{fl/fl}$ and FicD$^{-/-}$ mice.***
**Table S1**. Primer sequences for the genes analyzed in this study

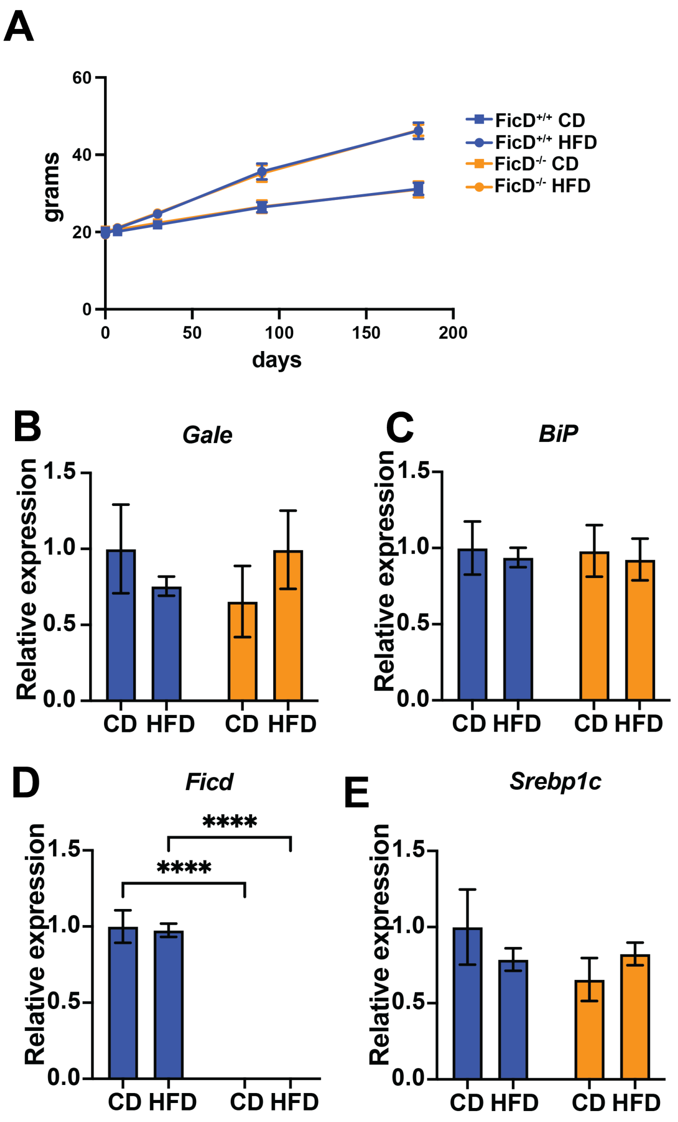


**Figure S1.** ***Gene expression in HFD feeding in FicD*$^{fl/fl}$ *and FicD*$^{-/-}$ *livers.*** A) Body weight measurements of *FicD*$^{+/+}$ *UMAI*$^{+/-}$ (blue) and *FicD*$^{-/-}$ *UMAI*$^{+/-}$ (orange) fed control diet (squares) and high fat diet (circles) over 6 months. B-E) Quantification of *Gale*, *Bip*, *Ficd*, and *Srebp1c* mRNA analyzed by qPCR from *FicD*$^{+/+}$ *UMAI*$^{+/-}$ (blue) and *FicD*$^{-/-}$ *UMAI*$^{+/-}$ (orange) mouse liver after6 months of CD and HFD feeding. Expression values were normalized to that of the housekeeping gene *U36B4.* Bars indicate mean relative expression compared to *Fic*$^{+/+}$ *UMAI*$^{+/-}$ CD controls, and error bars represent standard error. Statistics were performed using GraphPad Prism 9 using an 2-way ANOVA. N=5-6. *, $p < 0.05$; **, $p < 0.01$, ***, $p < 0.001$, ****, $p < 0.0001$.

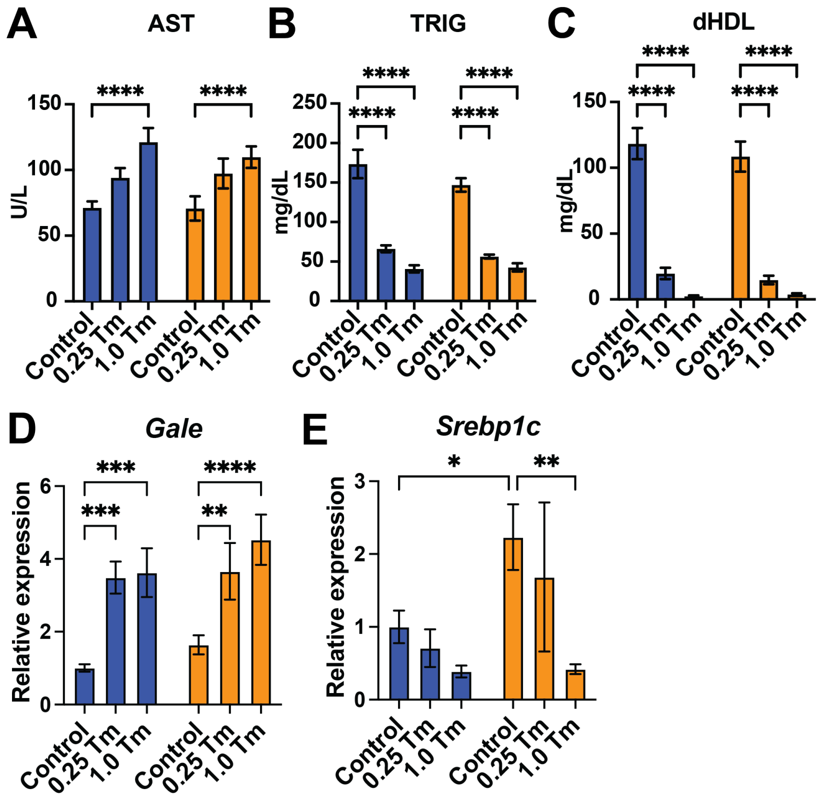


**Figure S2.** ***Hepatic metabolism regulation in tunicamycin treated FicD$^{fl/fl}$ and FicD$^{-/-}$ liver.*** A-C) Quantification of serum A) aspartate aminotransferase (AST) B) Total Triglycerides (TRIG), and C) high density lipoprotein cholesterol (dHDL) in Ficfl/fl and Fic-/- mice treated 24hrs with vehicle (control), 0.25mg/kg Tunicamycin, or 1mg/kg Tunicamycin. D-E) Quantification of *Gale* and *Srebp1c* mRNA analyzed by qPCR from *FicD*$^{fl/fl}$ (blue bar) and *FicD*$^{-/-}$ (orange bar) mouse liver 4 hours treatment with vehicle (control), 0.25mg/kg Tunicamycin, or 1mg/kg Tunicamycin. Expression values were normalized to that of the housekeeping gene *Gapdh.* Bars indicate mean relative expression compared to fasted controls, and error bars represent standard error. Statistics were performed using GraphPad Prism 9 using an 2-way ANOVA. N=5-6. *, $p < 0.05$; **, $p < 0.01$, ***, $p < 0.001$, ****, $p < 0.0001$.

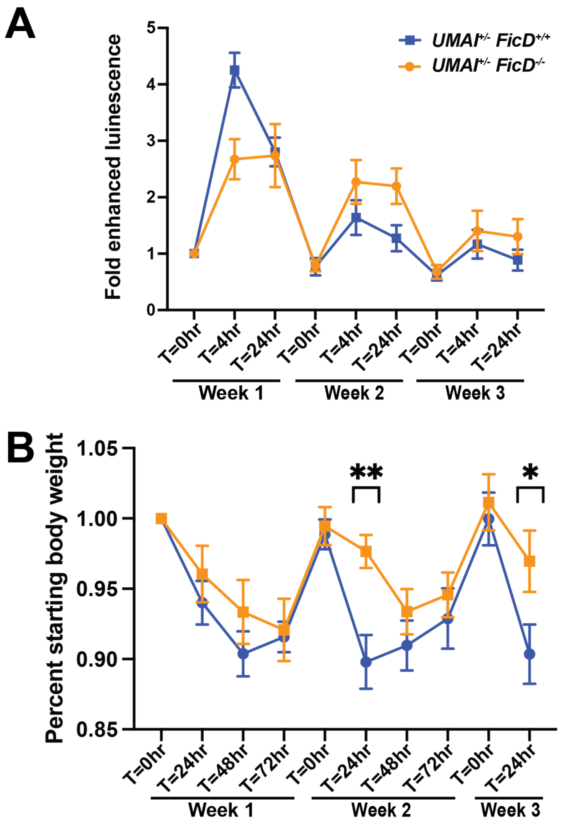


**Figure S3. Changes *during repetitive tunicamycin challenge and recovery in FicD$^{fl/fl}$ and FicD$^{-/-}$ mice.*** A) Fold induction of total body luciferase activity of *Fic$^{+/+}$ UMAI$^{+/-}$* (blue) and *FicD$^{-/-}$ UMAI$^{+/-}$* during three weekly rounds of 0.25mg/kg tunicamycin treatment. Measurements taken at T=0, T=4, and T=24hr after injection. Bars indicate mean relative expression compared to Week 1 T=0hr timepoint. Error bars represent standard error. Statistics were performed using GraphPad Prism 9 using an 2-way ANOVA. B) Normalized body weight of *FicD$^{+/+}$ UMAI$^{+/-}$* (blue) and *FicD$^{-/-}$ UMAI$^{+/-}$* (orange) during three weekly rounds of 0.25mg/kg tunicamycin treatment. Measurements taken at T=0, T=24, T=48hr, and T=72hr after injections. Symbols indicate mean change in body weight compared to Week 1 T=0hr timepoint. Error bars represent standard error. Statistics were performed using GraphPad Prism 9 using an 2-way ANOVA. **, $p < 0.01$. N=9

**Table S1**. Primer sequences for the genes analyzed in this study

| Gene name | Forward primer sequence | Reverse primer sequence |
|---|---|---|
| *Atf3* | 5’ tggagatgtcagtcaccaagtct 3’ | 5’ gcagcagcaattttatttctttct 3’ |
| *Atf4* | 5’ actctaatccctccatgtgtaaagg 3’ | 5’ caggtaggactctgggctcat 3’ |
| *BiP* | 5’ caaggattgaaattgagtccttctt 3’ | 5’ ggtccatgttcagctcttcaaa 3’ |
| *Chop* | 5’ ccagaaggaagtgcatcttca 3’ | 5’ actgcacgtggaccaggtt 3’ |
| *Fgf21* | 5’ cctctaggtttctttgccaacag 3’ | 5’ aagctgcaggcctcaggat 3’ |
| *Ficd* | 5’ gtagacgcactgaatgagttcg 3’ | 5’ tggtgtataagtagtcagcctgg 3’ |
| *Gale* | 5’ ccataacgccattcgtggag 3’ | 5’ tccagaggcttctgcactg 3’ |
| *Gapdh* | 5’ aggtcggtgtgaacggatttg 3’ | 5’ tgtagaccatgtagttgaggtca 3’ |
| *Pck1* | 5’ ttgaactgacagactcgccct 3’ | 5’ tgcccatccgagtcatga 3’ |
| *Srebp1c* | 5’ ggagccatggattgcacatt 3’ | 5’ ggcccgggaagtcactgt 3’ |
| *U36B4* | 5’ cgtcctcgttggagtgaca 3’ | 5’ cggtgcgtcagggattg 3’ |
| *Xbp1S* | 5’ctgagtccgcagcaggt 3’ | 5’ tgtcagagtccatgggaaga 3’ |
